# Whole genome sequencing of nearly isogeneic WMI and WLI inbred rats identifies genes potentially involved in depression

**DOI:** 10.1101/2020.12.04.411769

**Authors:** Tristan de Jong, Panjun Kim, Victor Guryev, Megan Mulligan, Robert W Williams, Eva E Redei, Hao Chen

## Abstract

**Background:** The WMI and WLI inbred rat substrains were generated from the stress-prone, and not yet fully inbred, Wistar Kyoto (WKY) strain using bi-directional selection for immobility in the forced swim test followed by over 38 generations of inbreeding. Despite the low level of genetic diversity among WKY progenitors, the WMI substrain is more vulnerable to stress relative to its WLI control substrain. Here we quantify numbers and classes of sequence variants distinguishing these substrains and test the hypothesis that they are nearly isogenic.

**Results:** The WLI and WMI genomic DNA were sequenced using Illumina xTen, IonTorrent and 10X Chromium technologies to obtain a combined coverage of over 100X. We identified 4,296 high quality homozygous SNPs and indels that differ between the WMI and WLI substrains. Gene ontology analysis of these variants showed an enrichment for neurogenesis related pathways. In addition, high impact variations were detected in genes previously implicated in depression (e.g. *Gnat2*), depression-like behavior (e.g. *Prlr, Nlrp1a*), other psychiatric disease (e.g. *Pou6f2, Kdm5a, Reep3, Wdfy3*) or stress response (e.g. *Pigr*).

**Conclusions:** The high coverage sequencing data confirms the near isogenic nature of the two substrains, which combined with the variants detected can lead to the identification of genetic factors underlying greater susceptibility for depression, stress reactivity, and addiction.

## Background

Major depressive disorder (MDD) is a common, debilitating disease that is the leading cause of “years lived with disability” worldwide [1]. Genetic factors play important roles in the etiology of MDD. Heritability of MDD is estimated to be between 28 and 44% [2,3], although recent estimates are over 50% [4]. Genomic variants contributing to depression have been difficult to identify, but large genome-wide association studies (GWAS) [5] are starting to identify candidates, including variants near *SIRT1, LHPP* [6], *OLFM4, MEF2C*, and *TMEM161B* [7]. Meta-analysis of GWAS based on self-reported depression also identified a larger number of independent and significant loci [8,9], although relying on self diagnosis may have reduced the reproducibility of the findings [6]. Even when the MDD diagnosis is not based on self-report, the current diagnostic methods are still comparatively subjective and cannot truly characterize subgroups of this complex disease, which are likely affected by differences in genetics. Thus, identification of sequence variants associated with the disease, and the genetic etiology of MDD, remains largely unsolved.

Compared to the high levels of genetic variations among humans (6 million between any two individuals), well defined animal models can tightly constrain both genomic and environmental variables. Many genetic mapping strategies have been developed for model organisms. For example, the reduced complexity cross (RCC) uses offspring from two genetically similar parents that have divergent phenotypes. The number of segregating variants in an RCC is orders of magnitude smaller than in conventional cross. It therefore often permits immediate identification of causal variants [10].

In this report we analyze a genetic model of depression and its nearly isogenic control strain, both bred from Wistar Kyoto (WKY) rats. The WKY strain had been developed as the normotensive control for the spontaneously hypertensive rat strain and was distributed to vendors and universities between the 4th and 11th generation of inbreeding [11]. At this early stage, the stock varied widely in behaviors [12]. The Redei lab obtained WKY rats from Harlan Laboratories (Madison, WI), where they had been bred for 65 generations. However, it is not known whether the sublines (breeding pairs) Harlan obtained at the beginning of the breeding were maintained as sublines or interbred. The WKY strain has become a well-established model of adult and adolescent depression and comorbid anxiety [13–15]. Its behavior mirrors several symptoms of human MDD and anxiety, including anhedonia, disturbed sleep, a reduced appetite and reduced energy, and the attenuation of depression-like behaviors after treatment with antidepressants [16–19].

A large variability in behavioral and psychological measurements were noted within the WKY strain [20,21]. The variability of behavior in the forced swim test (FST)—one of the most widely utilized tests for depressive behavior in rodents—motivated the bi-directional selection of the animals based on their level of immobility in the FST [22]. Males and females with the least mobility and lowest climbing scores in the FST were mated, producing the WKY *More Immobile* (WMI) line. Males and females with the highest mobility and highest climbing scores were mated, producing the WKY *Less Immobile* (WLI) line. Those animals showing the most extreme FST behavior within each line were selected for subsequent breeding, specifically avoiding sibling mating until the fifth G generation, when filial F matings were initiated.

Throughout the generations, the WMIs consistently have shown significantly greater immobility behavior in the FST than the WLIs [23]. The sex differences observed in the developmental pattern of MDD and its comorbidity with anxiety parallel differences observed in humans [24]. Maternal characteristics of the WMI after birth show similarities to that of women with postpartum depression [25]. Antidepressant treatments, specifically the tricyclic desipramine and the MAO inhibitor phenelzine, but not fluoxetine, alleviate depression-like behavior of WMIs [22], and enriched environment in adulthood does the same [26]. Resting state functional connectivity differences between WMIs and WLIs, measured by fMRI, are similar to those found in depressed patients [27,28]. Behavioral and hormonal responsiveness to acute and chronic stress also differ between the strains [26,29]. In humans, posttraumatic stress disorder (PTSD) and alcohol use disorder have high comorbidity with major depression. As hypothesized, the stress-reactive WMI strain showed increased fear memory in a model of PTSD, the stress-enhanced fear learning behavior compared to the isogenic WLI strain [29]. Additionally, WMIs consume more alcohol than WLIs when tested using an operant licking procedure [29]. In human studies depression has been noted as a risk factor for dementia in females. Similarly, middle-aged WMI females show cognitive decline compared to middle-aged WLI females [30]. Together, these data establish WMI as a suitable model to study human depression.

The WMI and WLI strains also differ in their brain and blood gene expression profiles [23]. A panel of blood transcriptomic markers, developed using the WMI strain, can diagnose major depression in humans. These blood transcriptomic markers are able to distinguish adolescent and adult subjects with major depression from those with no disorder with a high level of reliability [31–33]. Additionally, the expression of these markers correlated with depression symptoms in pregnant women [34]. These data provide tantalizing evidence that genetically determined gene expression differences between the WMI and WLI substrains can potentially lead to the discoveries of molecular mechanisms of depression in humans.

Full genome sequencing provides an abundance of genetic information (single nucleotide polymorphisms, inserts and deletions, and large structural variants) and can allow for comparative genomics between the rat model and humans. Comparing the genome of WMI and WLI to each other as well as the reference genome could provide insights to the underpinnings of their distinctive behavioral phenotypes. Because the WMI and WLI strains were both derived from WKY founders, we hypothesized that a small number of genetic variants between these strains contribute to behavioral and physiological differences in depression-associated traits between WMI and WLI. Here we describe the whole genome sequencing of these two strains using data obtained from three different platforms (Illumina xTen, Ion Proton, and 10X Chromium linked-read) and the identification of genetic variants between these strains.

## Results

To discover SNPs associated with the depression phenotype in WLI and WMI rats, whole genome sequencing data was obtained using three different platforms: Ion Torrent Proton, 10X Chromium and Illumina xTen from male WLI and WMI rats. Each technique covered an average depth of 41, 27 and 43 for both strains, respectively (Figure 1). X-chromosomal coverage was expected to be half of autosomal coverage but was found to be much higher on IonProton and Illumina X-ten sequencing results (Figure 1).

**Figure 1.**
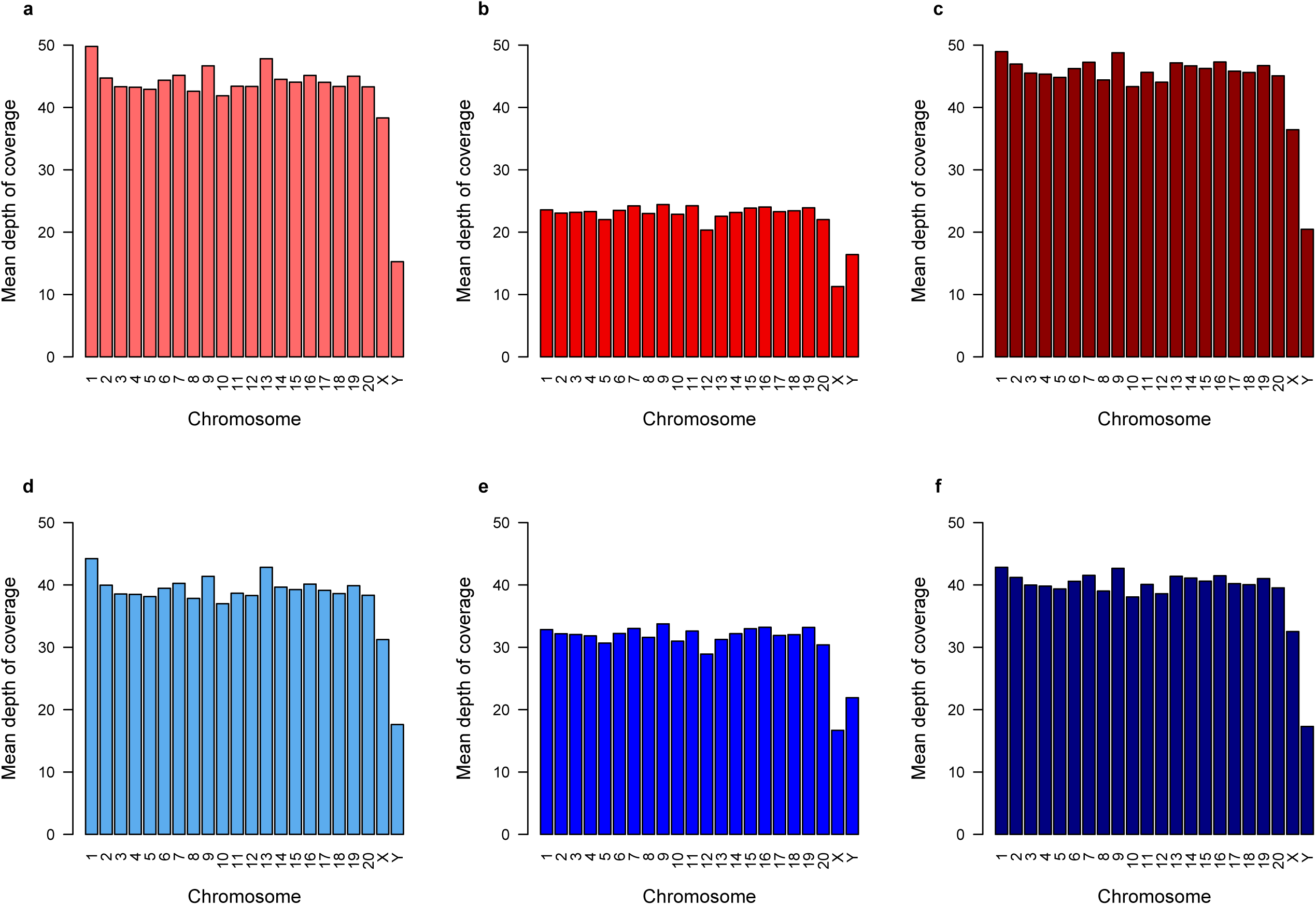
Mean depth of coverage per chromosome per sample. Deepvariant called a total of 12,764,518 unique variants across 20 chromosomes plus X and Y with varying quality scores on either WLI or WMI samples. Depth of coverage is shown per A) WLI, IonProton, B) WLI 10X Chromium, C) WLI Illumina xTen, D) WMI IonProton, E) WMI 10X Chromium, F) WMI Illumina xTen.

Sequencing data were mapped to the rat reference genome rn6 using bwa (Illumina and IonProton data) or LongRanger (10X Chromium data). The resulting bam files were used as the input to DeepVariant to report genomic variants (i.e. SNPs and small indels) for each sample. GLNexus was then used to conduct a joint analysis of variants across all six samples. Over 12 million unique variants were identified before filtering. The analysis workflow was designed to take full advantage of the data provided from three sequencing technologies. We are interested in variants that have a Phred quality score above 30, have a clear call for either reference, homozygous or heterozygous, have no matching calls between WLI and WMI and must not have both a high quality reference and alternative allele called on different sequencing platforms within the same strain (Figure 2).

**Figure 2.**
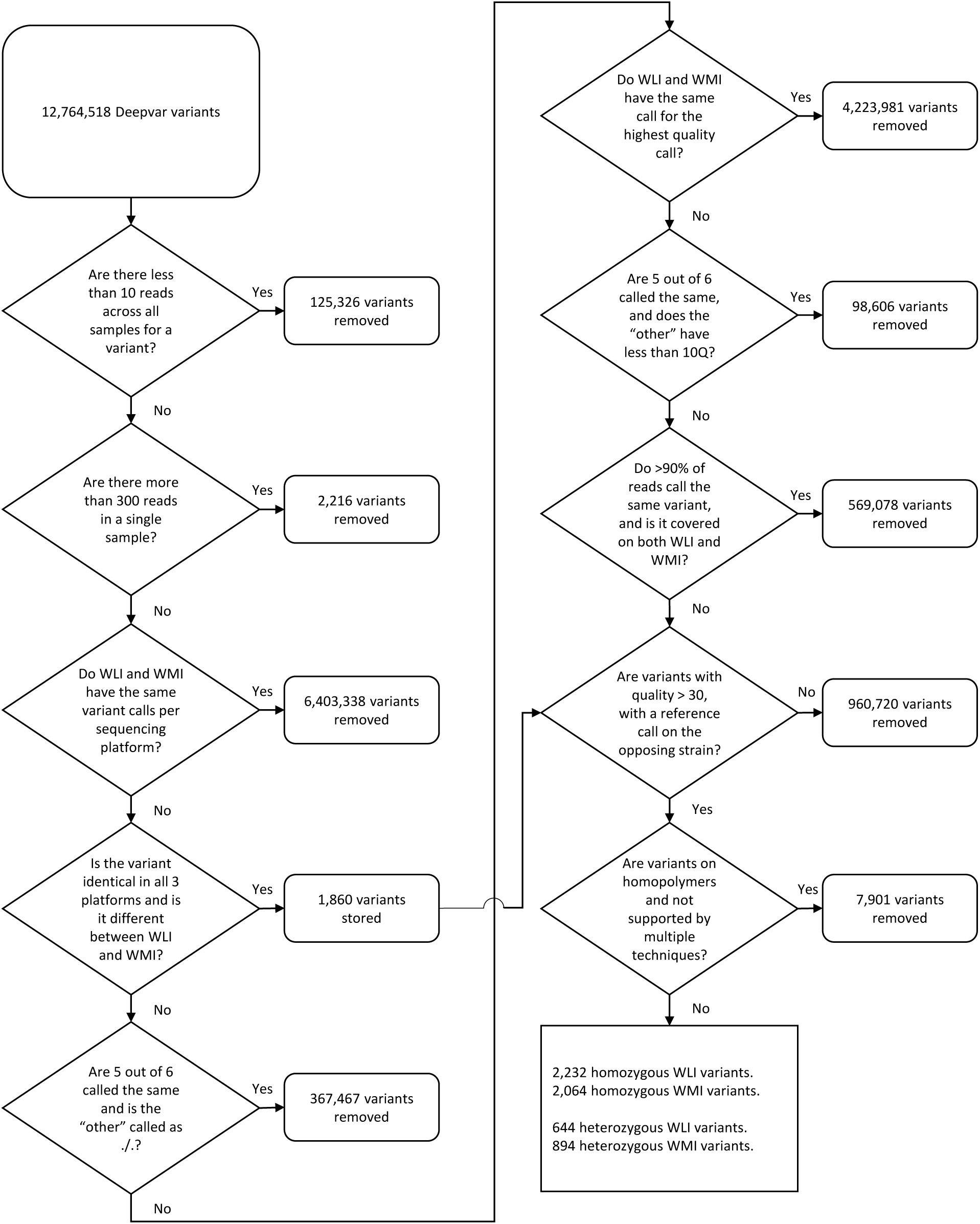
Flowchart of each filtering step and the number of variants removed per step. The initial 8 steps were performed in Python, the last 2 were performed in R.

A large portion of the variants had a Phred quality score below 10 and were excluded from subsequent analysis. In total, 99,465, 25,937, and 6,454 homozygous variants had a Phred quality score greater than 10, 20, and 30 in at least one sample, respectively. For heterozygous calls the number of variants were ∼3 million, ∼1 million and ∼200 thousand, for quality scores of 10, 20 and 30 respectively. The number of high-quality calls for homozygous variants varied per sequencing technology (Figure 3).

**Figure 3.**
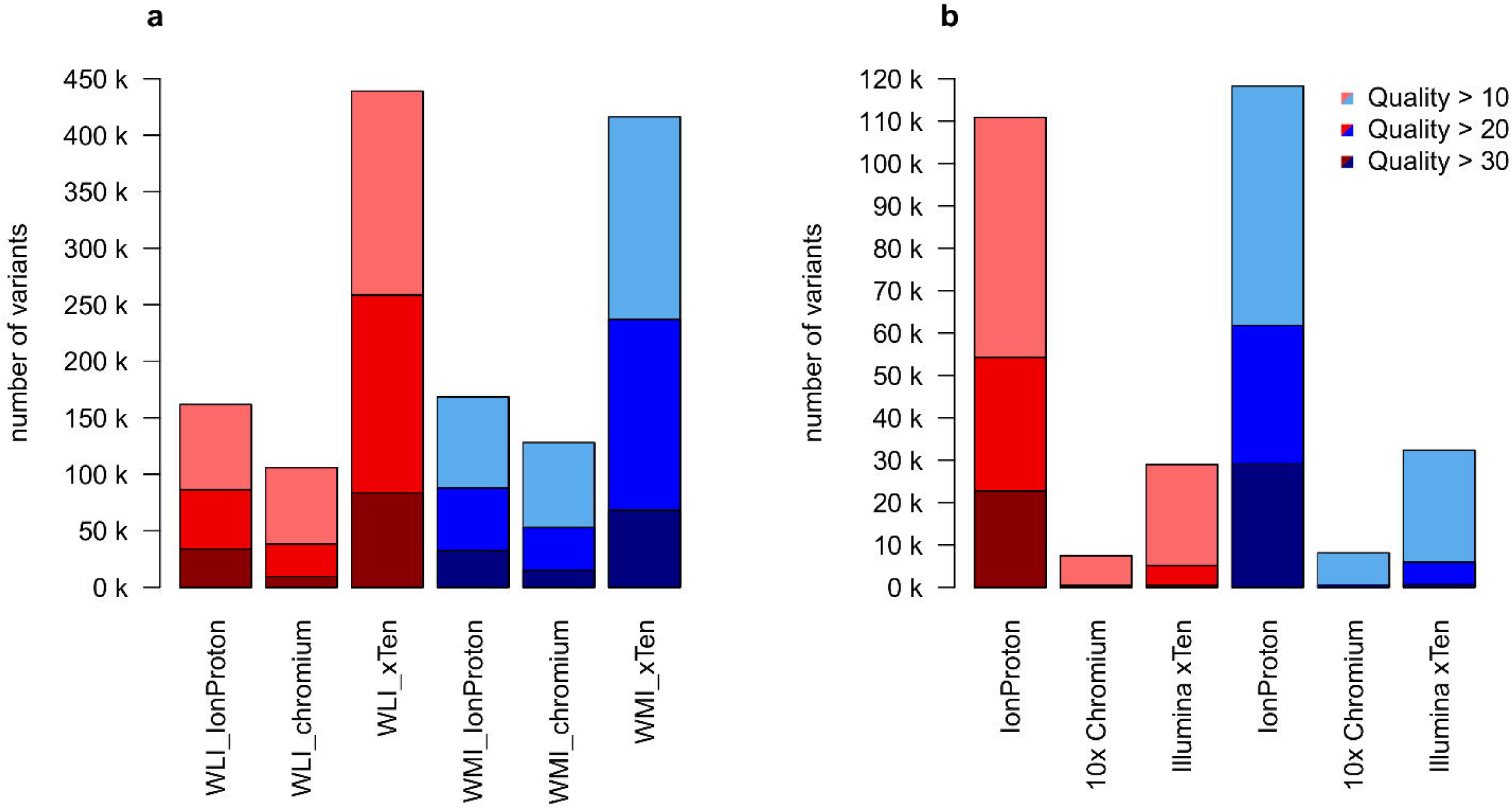
A) Number of ALT calls by deepVariant. Samples are separated based on quality score: quality 10 = (p < 0.1), quality 20 = (p < 0.01), quality 30 = (p < 0.001). B) Number of HET calls by deepVariant in different samples separated by quality.

**Figure 4.**
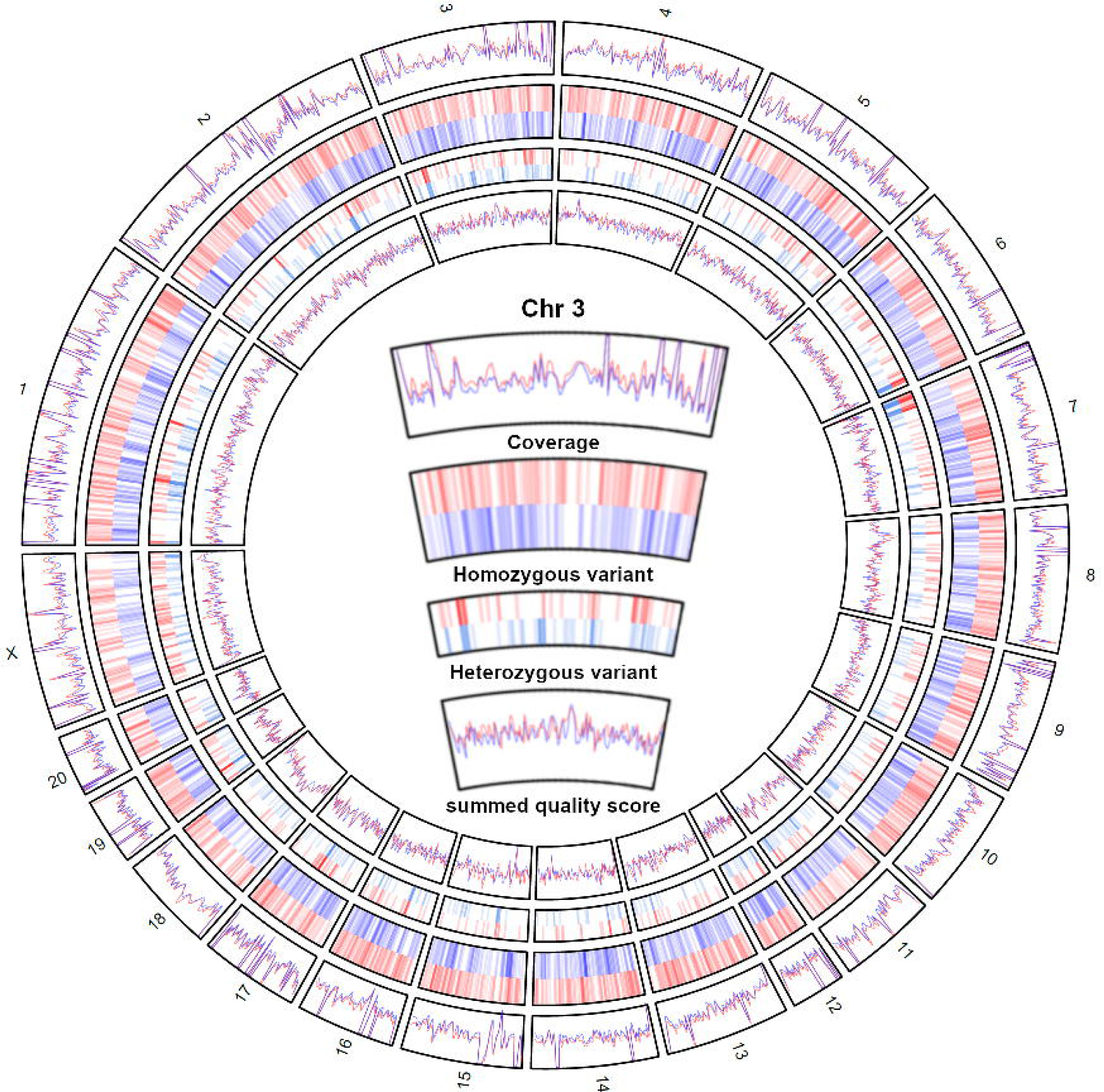
From outside to inside: 1) Smoothed summed coverage of variant calls per technique for WMI samples (blue) and WLI samples (red). 2) Hotspots of homozygous SNPs on each chromosome found only in WMI (Blue) or WLI (Red). 3) Hotspots of heterozygous variants on each chromosome found only in WMI (light blue) or WLI (light red). 4) Smoothed summed quality of variant calls per technique for WMI samples (blue) and WLI samples (red).

The majority of high confidence heterozygous calls came from a single technique, Ion proton (Figure 3). Closer inspection revealed that the majority (>90%) of these calls was detected on homopolymeric nucleotide sequences. In addition, approximately 95% of these were deletions rather than SNPs, further confirming that these calls are due to errors in base calling homopolymeric sequences. To filter out this common sequencing error, all deletions on homopolymeric regions which were not supported by at least one other sequencing technique were removed (Supplementary figure 1).

As a final result, 2,232 and 2,064 homozygous high confidence variants were discovered on WLI and WMI respectively (Table 1). The majority were insertions (45.3%) followed by SNPs (36.9%), and finally deletions (17.8%) (Table 1). Of these SNPs, approximately 57% were transitions, meaning a purine nucleotide was mutated to another purine or a pyrimidine nucleotide to another pyrimidine. The other 43% were transversion SNPs, in which a purine was replaced by a pyrimidine or vice versa (Table 1). A total of 655 and 894 heterozygous variants were identified for WLI and WMI. It should be noted that the heterozygous variants contained higher coverage than average as compared to homozygous variants (supplemental Figure 2). This implies a large portion of these could be homozygous SNPs aligned to collapsed regions on the reference genome.

**Table 1.** Overview of the number of variants, insertions and deletions in the final selection per strain.

|  | **WLI** | **WMI** |
| --- | --- | --- |
| **Transition SNP** | 478 | 428 |
| **Transversion SNP** | 325 | 356 |
| **Insertions** | 1090 | 855 |
| **Deletions** | 339 | 425 |

Cross comparison of the genomic positions of variants discovered on WLI and WMI with variants discovered in a panel of 44 inbred rat strains (our unpublished data) allowed us to gain an indication into whether the variants were likely de novo. Out of the 2232 variants on WLI and the 2,064 variants on WMI, 1215 and 856 were unique among all strains, respectively.

In addition, 79 and 119 homozygous variants were identified for WLI and WMI, respectively, with a Phred quality score of at least 10 in all three sequencing technologies. Though verified across technologies, quality scores cannot be simply summed. For certitude these were not included in the final selection.

We used SnpEff [35] to identify the impact, location (table 2), and the nearest gene in proximity of these variants. About half of the variants (52%) are located within intergenic regions, whilst some (a total of 62) variants fall within exons, 2,432 are within introns, and 450 are located within 5 KB upstream of a gene (Supplementary table 1).

**Table 2.**
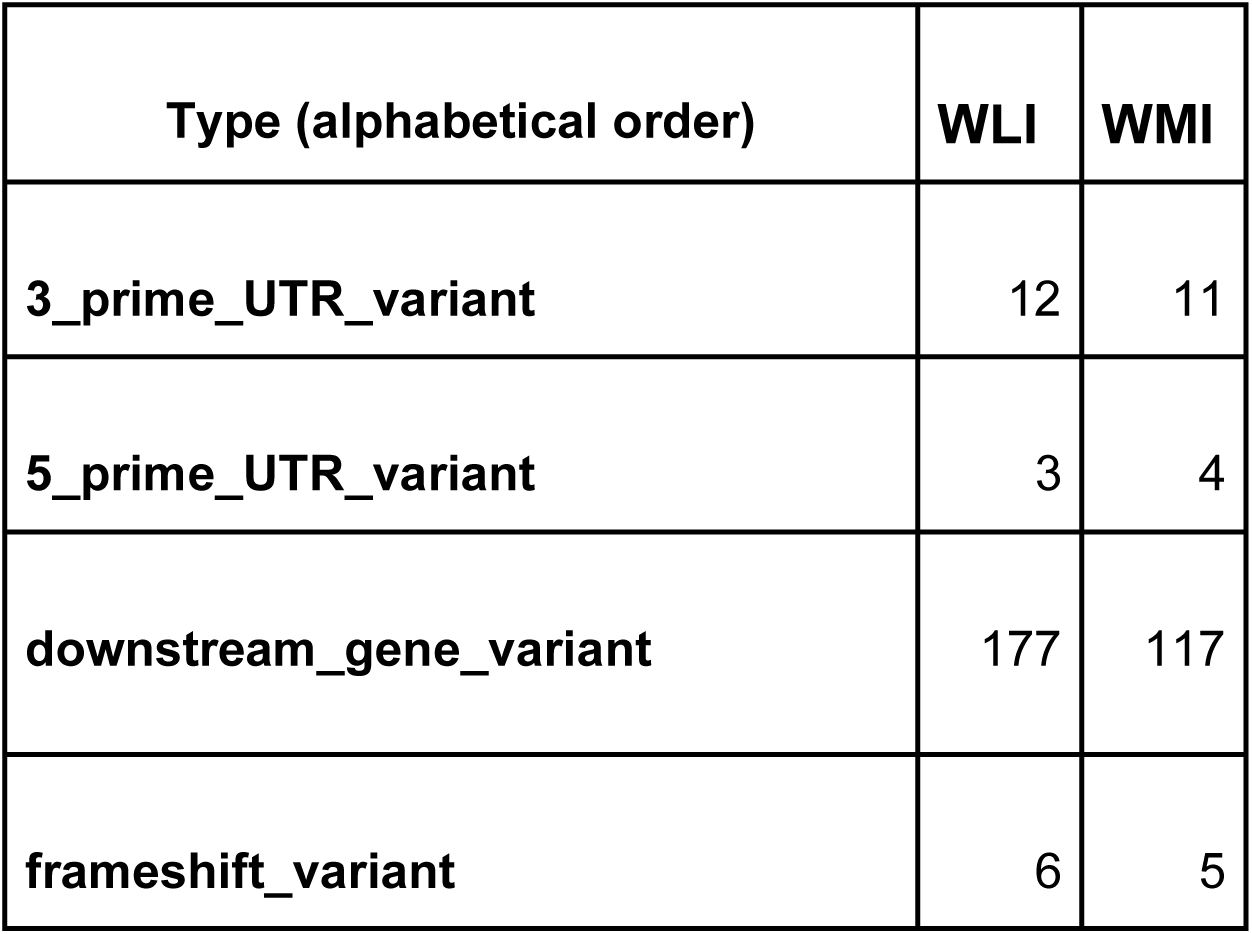
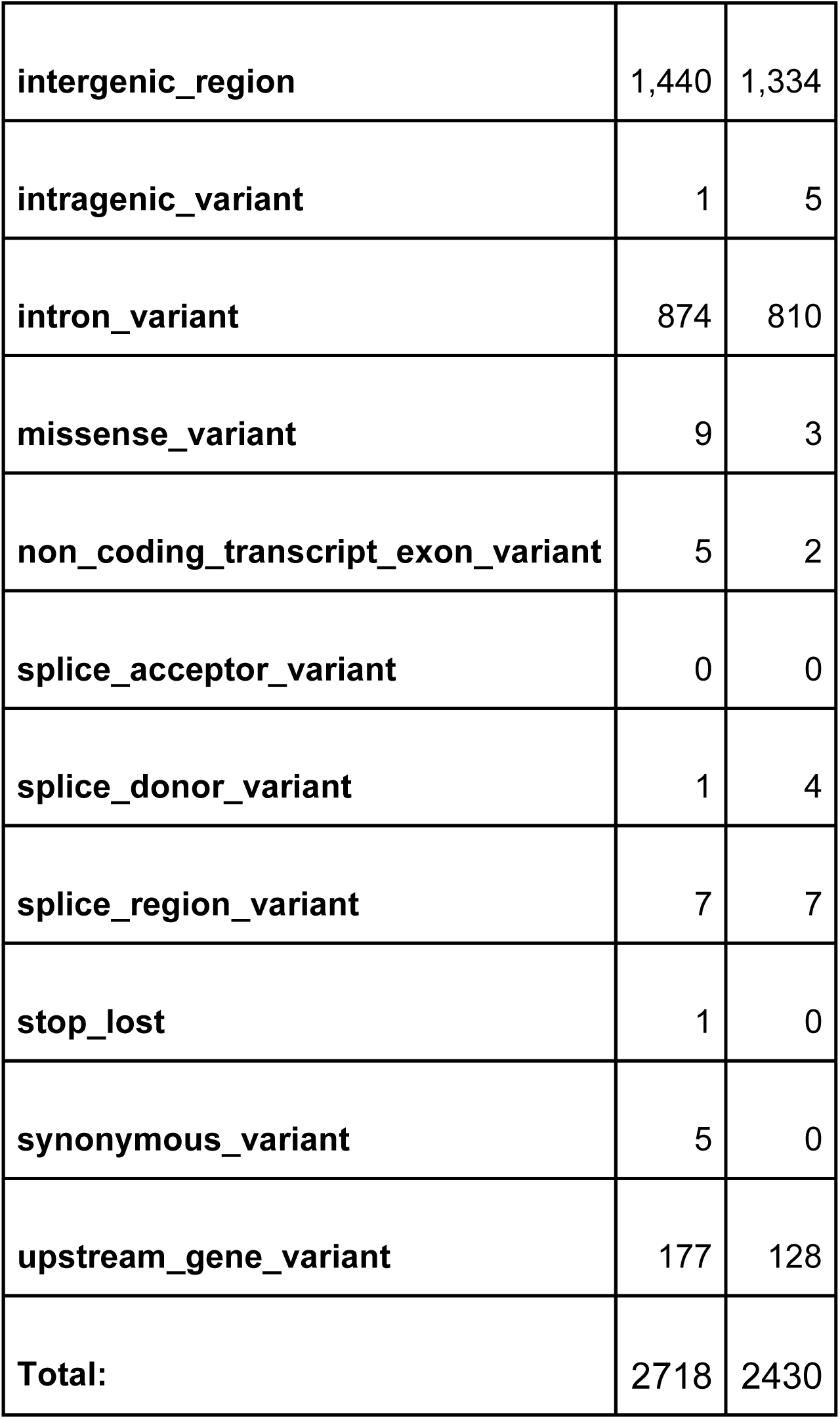
Position of selected variants in regions of interest.

| **Type (alphabetical order)** | **WLI** | **WMI** |
| --- | --- | --- |
| **3_prime_UTR_variant** | 12 | 11 |
| **5_prime_UTR_variant** | 3 | 4 |
| **downstream_gene_variant** | 177 | 117 |
| **frameshift_variant** | 6 | 5 |
| **intergenic_region** | 1,440 | 1,334 |
| **intragenic_variant** | 1 | 5 |
| **intron_variant** | 874 | 810 |
| **missense_variant** | 9 | 3 |
| **non_coding_transcript_exon_variant** | 5 | 2 |
| **splice_acceptor_variant** | 0 | 0 |
| **splice_donor_variant** | 1 | 4 |
| **splice_region_variant** | 7 | 7 |
| **stop_lost** | 1 | 0 |
| **synonymous_variant** | 5 | 0 |
| **upstream_gene_variant** | 177 | 128 |
| **Total:** | 2718 | 2430 |

In total, 1491 unique genes were in closest proximity to the final selection of homozygous variants across both strains. Of these, 744 genes were found in WMI and 866 in WLI (119 were found in both strains).

In total, 9 WMI variants and 11 WLI variants were estimated to have a large impact on the final protein product. These included changes to splice sites, missense mutations, loss of stop codons or frameshifts (Table 3). These genes included *Asxl1, Zfp292, Wrap73, Col5a3, Abcc5, Fscn1, Wdfy3, Pou6f2, Svil, Prlr, Gnat2, Slc30a7, Kdm5a, Slco1a2, Nlrp1a, Crlf3, Tpcn1, Pigr, Pou6f2* and *Reep3*. Of these, five variants are likely de novo, whilst 15 are also found on other strains. (Table 3). Among these genes, *GNAT*2 has a variant in human (rs6537837) that was reported to be associated with unipolar depression with a genome wide significance of p=1e-6 [36], while *Prlr* and *Nlrp1a* were implicated in depression-like behavior in animal models [37,38]. Further, *Pou6f2, Kdm5a, Reep3, Wdfy3* have been implicated in psychiatric diseases such as autism [39–41] and *Pigr* was found to be involved in stress response [42].

**Table 3.**
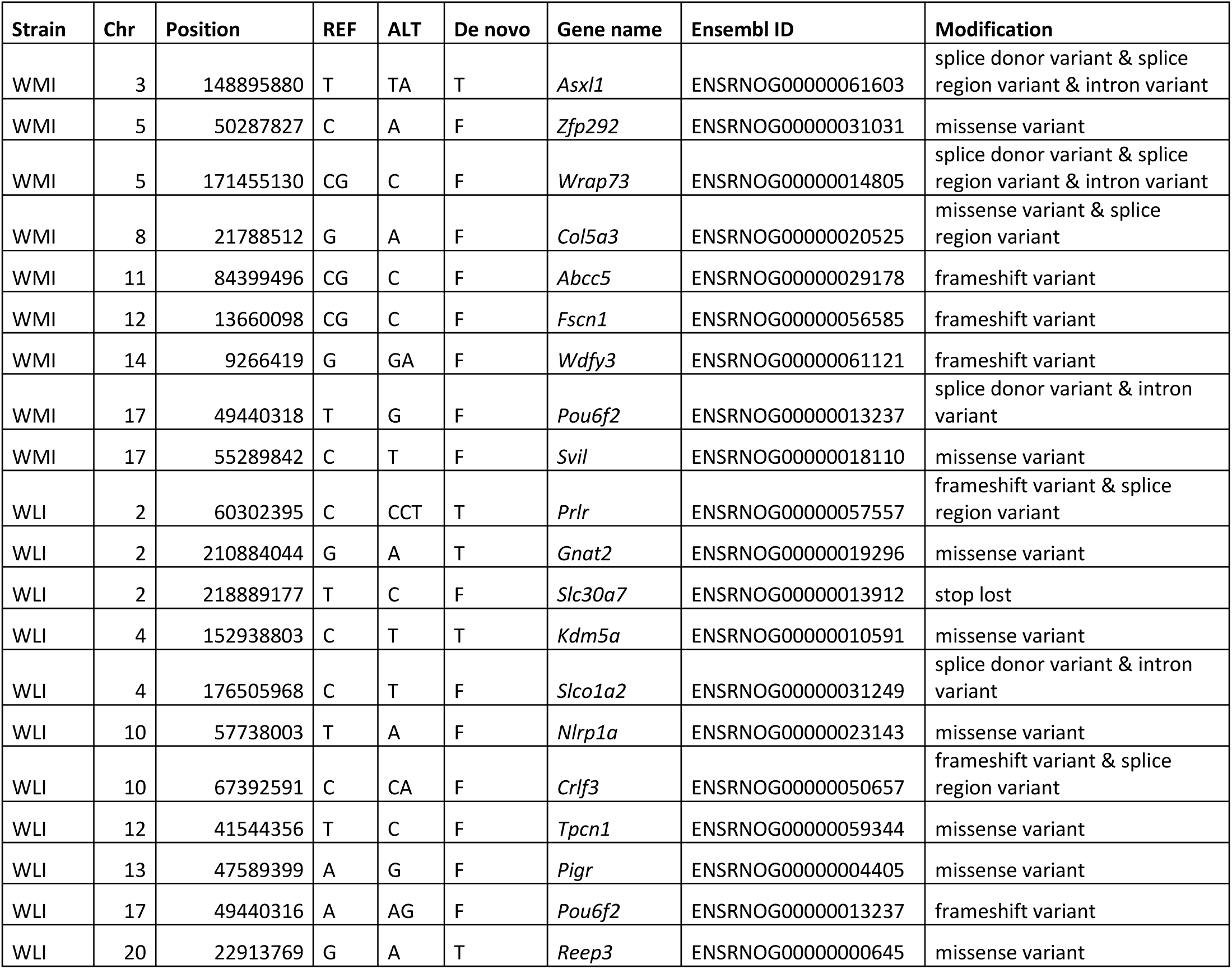
Overview of variants of high and moderate impact, likely de novo status, their impact and the gene affected.

We also leveraged Gene Ontology-term (GO-term) and KEGG-term enrichment analysis using G-profiler (https://biit.cs.ut.ee/gprofiler/gost) to explore the biological functions of genes in close proximity to sequence variants. We found an over-representation of several neurogenesis, behavioral and locomotion related pathways. Over-represented terms for WLI included locomotory behavior, behavior, nervous system development, neuron projection and neurogenesis (supplementary table 2a). Over-represented terms for WMI included Par-3-KIF3A-PKC-zeta complex, actin-mediated cell contraction, neuronal related and cellular stress related pathways (supplementary table 2b).

Of the genes found in close proximity to high impact variants (based on SNPeff annotations) in WLI, 30 were annotated with the GO-term neuron to neuron synapse (GO:0098984). We further examined these genes using RatsPub [43], an online tool that allows us to conduct automated searches for genes associated with depression, addiction, stress, or other psychological afflictions from PubMed. We found 23 genes that were associated with psychiatric disease in previous research. These genes included; *Syt1, Stxbp5, Sorcs2, Rs1, Ptprd, Prkcz, Pdlim5, Lyn, Itga8, Igsf11, Grm3, Erbb4, Epha7, Epha4, Dlgap1, Dgki, Cdkl5, Cacna1c, Atp2b2, Ank2, Als2, Add3, Adcy8* (Supplementary table 3).

Lastly, we validated our genome sequencing results by selecting 224 SNPs, half unique to each strain, and using multiplex PCR to amplify each target region (150 bp flanking each variant) in genomic DNA collected from 8 rats, including four WMI and four WLI with equal number of males and females. We then constructed sequencing libraries using these PCR products and sequenced them on an Illumina instrument. We were able to obtain PCR products from 89 and 87 primers sets targeting WLI and WMI specific variants, respectively. Among them, 75 WLI and 76 WMI targets met the following two criteria: 1. homozygous alternative in at least three rats of the target strain; 2. homozygous alternative in none of the rats of the opposite strain. Therefore, the positive rate of our stringent empirical validation using 8 rats was 85.8%.

## Discussion

The goal of this research is to give us genetic markers for WLI and WMI in context of other strains in reduced complexity crosses and to give us candidate variants for immediate scrutiny of linkage to depression. We used three leading next-generation sequencing technologies to obtain a combined coverage of approximately 100X for each genome of two closely related inbred rat strains, the WMI and WLI. We identified 4,296 homozygous variants with high fidelity that are located in close proximity to 1,491 unique genes that differ between these two strains. The SNPs and indels identified in this dataset offer new opportunities for the identification of genes related to the phenotypic differences between the WLI and WMI strains.

Each of the three sequencing methods we used has its own merits and flaws. For example, compared to the widely used Illumina platform, the Ion Torrent platform provides high quality data at a lower cost. However, it suffers at homopolymer regions. The 10x Chromium linked reads technology attaches barcodes to high molecular weight DNA before library preparation and can detect large structural variants. But obtaining good quality HMW DNA is technically challenging and is associated with increased cost. Further, when utilizing sequencing data from a single technique, technical biases are likely to make their way into the final result. By removing variants called differently by sequencing platforms, the technical bias is mitigated across the final selection of variants.

We used DeepVariant to identify SNP and small indels across all sequencing techniques [44]. Deepvariant has been shown to outperform GATK in different tests, especially in calling indels [45]. It also handles data from diverse sequencing platforms without additional calibration. We also used LongRanger to map the 10X linked reads sequencing data to the reference genome because it incorporates the molecular barcodes into the mapping algorithm. Following DeepVariant analysis, we used GLNexus to conduct a joint analysis to obtain a raw list of genomic variants. Joint analysis empowers variant discovery by leveraging population-wide information from a cohort of multiple samples, allowing us to detect variants with great sensitivity and genotype samples as accurately as possible [46].

With the combination of different sequencing methods, a higher certainty of variant calling between WLI and WMI has been made possible, though throughout this experiment strict filtering was performed. We first removed any variants detected with any certainty in both WLI and WMI, because we are interested in the differences between these two strains, rather than the common differences to the reference genome. One caveat of this approach is that variants incorrectly called by DeepVariant (e.g. due to low coverage in a single method) can lead to the exclusion of potentially interesting targets. Similarly, a strict quality score cutoff of 30 was used whilst un-opposed by any other sample with a Phred quality score of 10 or higher. Based on the abundance of data we have collected, including thousands of identified variants, we decided that 1 in 1000 variants was a sensible cutoff to avoid hundreds of false positives in the final set of variants.

The current reference genome (rn6) consists of 75,697 contigs and 1,395 scaffolds with N50 lengths of 100.5 KB and 14.99 Mb respectively. These sequences combine into a golden path of approximately 2.8 billion bases. Due to the fragmented nature of the reference genome, the identification of structural variants has proven to be difficult. One example of this is that it is often not possible to establish whether sequence variation is strain specific or related to a problem with the reference genome. In addition to the 4,296 high quality homozygous variants discovered in this research, an additional 15,268 variants were discovered in either WLI or WMI with no significant coverage or significant phred score on the opposing strain (called as ./.). Without a high quality read on both strains we cannot verify newly discovered variants. These low quality calls could be caused by heterozygosity, low coverage or overlapping variants. Initial studies using a new rat reference genome (mRatBN7.1, yet to be annotated) has shown that low quality variant calls become much more sporadic with the updated reference genome (de Jong et al. Unpublished results). For this reason, we have opted to exclude these high quality variant calls without a quality call on the opposing strain.

In this investigation 655 and 894 heterozygous variants were discovered on WLI and WMI respectively. Despite both strains being fully inbred, there is a chance that de-novo mutations could propagate as heterozygous variants within each substrain. A look at the coverage of these positions reveals an average two-fold coverage, implying these variants are homozygous on collapsed regions on the reference genome, or duplicated and mutated within either strain. With an updated reference genome these regions could be resolved and can contribute in a meaningful way to the identification of variants that contribute to phenotypic differences between WLI and WMI substrains.

Ongoing research has identified over 40,000 variants in multiple BN/NHsdMcwi samples (the strain used to generate data for rn6). This means some caution is required when identifying variants for WLI or WMI strains based on comparison to the current rat reference genome. There is a chance that variants found in both strains could potentially be due to base level errors in the reference, i.e., there is no variant present at all. Similarly, when variant is only reported in strain A, there exists a small chance that the variant actually is located on strain B (i.e. the base level error in the reference happens to be the same as the sequence in strain B). Thus, a small percentage of the reported mutations in WLI strain could potentially be present in WMI. This might contribute, to some degree, to the enrichment of neuronal GO-term annotation for genes located within the vicinity of WLI sequence variants.

GO-term annotation enrichment for genes in the nearest proximity of variants detected in WLI included locomotor behavior and neuron projection. This provides some evidence that these variants could be capable of producing an impact on behavior, however this will require further investigation. As locomotory behavior is a complex trait, a combination of variants can be causal. For WMI the terms: neuron to neuron synapse (GO:0098984), nervous system development (GO:0007399), generation of neurons (GO:0048699), and finally, the Par-3-KIF3A-PKC-zeta complex (CORUM:899) was significantly over-represented. The Par-3-KIF3A-PKC-zeta complex is interesting as both parts are in proximity of variants detected on WMI and it is involved in the establishment of neuronal polarity [47].

The ancestral WKY strain was noted for its highly variable behavior [19,20]. The WLI and WMI have been selected for both depressive and non-depressive behavior. With the discovery of variants associated with psychiatric phenotypes in both strains it should be kept in mind that variants could have been both selected for and against. In addition, as discussed above, there is a small chance some variants are located on the opposite strain due to potential errors in the reference genome. For this reason, we have only included variants which are different between WLI and WMI and not those that are different relative to the reference genome. The WMI strain has been assigned to be a genetic model of depressive behavior, but the functional selection using the forced swim test could illuminate the potential connectedness of multiple phenotypic differences between the strains. The forced swim test is arguably thought of as a measure of stress-coping strategy [48], therefore, many behavioral phenotypes that employ stress coping could differ between the strains.

The small number of variants between the WMI and WLI strains indicates that they are close to isogenic. And yet, there exist numerous phenotypical differences between these two strains. This provides an opportunity to use genetic mapping strategies such as reduced complexity cross [10] to discover causal variants mediating behavioral phenotypes such as susceptibility to depression, stress reactivity, learning, memory, aging and drug abuse.

## Methods

### Animals

Liver tissue from 4 adult WLI (2 males and 2 females) and 4 adult WMI (2 males and 2 females) rats were collected. Equal amounts of tissue from males and females were pooled for each strain (total weight = 20 mg). DNA were extracted using the Qiagen DNeasy blood and tissue kit (Cat# 69506).

### Whole Genome Sequencing

For sequencing using the HiSeq X Ten instrument, DNA whole genome shotgun sequencing libraries were generated using 200 ng of genomic DNA as input for the TruSeq Nano DNA Library Prep Kit (Illumina). Indexed libraries were sequenced as pools of eight samples on a full slide (8 lanes) on an Illumina HiSeq X Ten sequencer using HiSeq X Ten v2.5 reagents. For sequencing using the Ion Torrent instrument, 1 μg of genomic DNA was sheared to an average size of 200 bp using a Covaris S2 Sonicator. Then 500 ng of the sheared DNA was used to prepare libraries for sequencing using the AB Library Builder(tm) Fragment library Kit on a Library Builder system. Libraries were used without amplification and size selected on a 2% Pippin Prep gel. After quantification using qPCR, the libraries (190 pg) were then used to prepare beads for sequencing using an Ion Torrent One Touch instrument. DNA on these beads then sequenced on an Ion Torrent Proton sequencer using Hi-Q chemistry and a P1 chip. For 10X Chromium sequencing, the Qiagen MagAttract HMW DNA kit was used for DNA isolation. Sequencing library was then constructed from 1 ng of high molecular weight (∼ 50kb) genomic DNA using the Chromium Genome Library kit and sequenced on Illumina Hi-Seq (150 bp PE).

### Mapping

Illumina and Ion proton data were mapped to the rat reference genome (rn6) using bwa (reference). 10x Chromium data were mapped to rn6 using LongRanger (ver 2.2.2). DeepVariant (ver 1.0.0) was used to call SNPs and small indels from the bam files and GLnexus was used for joint calling of variants.

### Analysis

Variant identification was performed separately for each strain and sequencing method. A total of 6 samples spread over 2 strains and 3 sequencing technologies were analyzed. Variants with less than 10 reads across all samples or more than 300 on a single sample for a variant were removed. Variants with the same highest quality call for WLI and WMI were removed. Variants with an identical call for all three sequencing technologies within either WLI or WMI were stored for further analysis. Variants with 5 out of 6 uncertain calls (./.) were removed. Variants with the same highest quality call for WMI and WLI were removed. Variants with 5 out of 6 identical calls of which the last had a quality score less than 10 were removed. If the majority (>90%) of reads were of the same variant call across all reads and both strains shared at least 25% of all reads, the variant was removed.

Variants were selected based on the highest quality call per method and removed if disputed by variants called on another sequencing method with call quality of at least 30 within the same strain. Only variants were included in which the call for WLI differed from WMI and one of two strains was called as 0/0 (reference allele). Finally, all deletions on a position consisting of two identical nucleotides (homopolymeric) which were not supported by multiple sequencing techniques were removed (Figure 2). The final selection was exported to VCF per strain and type of call (homozygous or heterozygous).

Variants were identified in a panel of 44 inbred rats samples (5 BN samples, 32 HXB recombinant inbred strains including parental strains and 7 other inbred strains, our unpublished data). The genomic positions of variants were cross-compared with the variants of WMI and WLI without specific filtering or pre-selection to identify the number of likely de novo variants on WMI and WLI.

SnpEff (v4_3t_core) was used for nearest gene identification, impact estimation and annotation of the VCF for selected variants [35]. Impact and nearest genes were estimated separately per strain, as well as heterozygous and homozygous variants. Variants marked as high or moderate impact were separated and placed in table 3. The annotated VCF is available for reference. g:Profiler version e101_eg48_p14_baf17f0 was used for GO-term enrichment analysis, standard settings were used, no background dataset was utilized [49]. RatsPub [43]) was used to explore a small set of genes nearest to variants enriched with the GO-term: neuron to neuron synapse (GO:0098984).

### Validation of variants and small indels by targeted re-sequencing

Ear punches from four WLI, four WMI (equal number of males and females) were used to extract genomic DNA. A total of 112 variants unique to WMI and 112 variants unique to WLI were selected from the final list of variants, with approximately equal distribution across the genome. Individual primer pairs were designed using Batch Primer 3 (http://probes.pw.usda.gov/batchprimer3/) at default settings for generic primers with total amplicon size set as an optimum of 100bp with the amplified region containing the target SNP (or region of interest). The primer sequences and genomic DNA were submitted to Floodlight Genomics (FG, Knoxville, TN) for processing using a Hi-Plex targeted sequencing approach [50]. The Hi-Plex approach pools primers to PCR amplify targets and adds a barcode sequence during the amplification process. The resulting target library is then sequenced on an Illumina instrument. Data were then aligned to the fasta file containing the targeting target variants using bwa. Genotypes for each sample were called using DeepVariant.

## Figure legends

**Supplementary figure 1.** Supplementary figure 1. Total number of homozygous (ALT) and heterozygous (HET) variants after final selection before and after homopolymer removal per strain.

**Supplementary figure 2.** Coverage of reference called variants to alternative on opposing strains (REF), homozygous variants (ALT) and heterozygous variants per sequencing technology. The coverage of heterozygous variants is overal twice as high as reference calls to homozygous variants on the opposing strain and homozygous variants.

**Supplementary table 1.** Positions of variants relative to genes on the genome for both WLI and WMI. Some regions overlap in classification.

**Supplementary table 2 A)**. GO-term enrichment analysis for WLI and **B)** WMI of selected homozygous variants.

**Supplementary table 3.** Overview of 23 out of 30 genes associated with the enriched GO-term: neuron to neuron synapse (GO:0098984) that have been associated with psychiatric disease in previous studies.

## Notes

### Competing Interest Statement

The authors have declared no competing interest.

### Summary of Updates

Genes are now in italics as they should be. Instead of mentioning shared genes, selected genes are stated between WLI, WMI and other strains. Clarification of goals added to text for overall readability.

