## Supplementary figures and images for "Whole genome sequencing of nearly isogeneic WMI and WLI inbred rats identifies genes potentially involved in depression"

### Supplemental Figure 1.

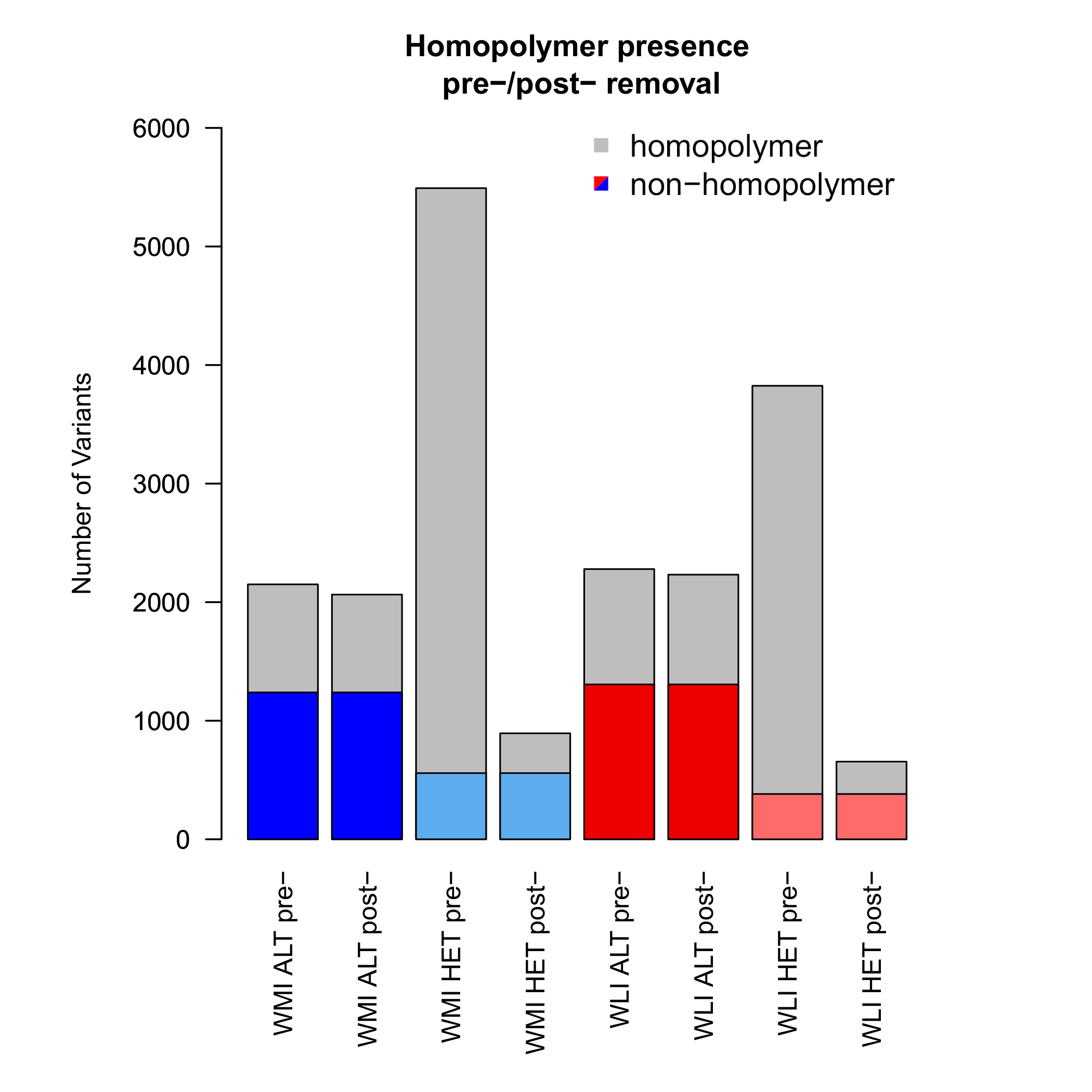

### Supplemental Figure 2.

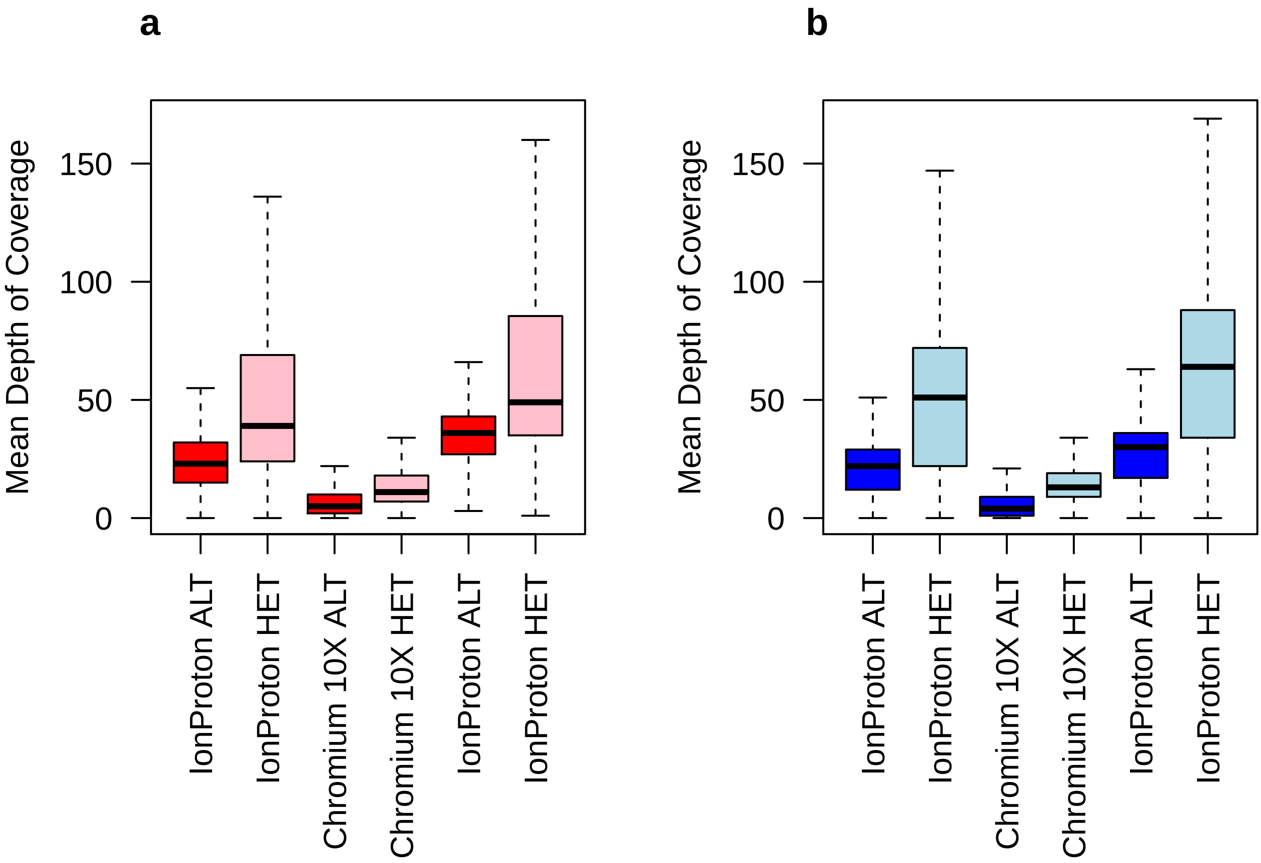

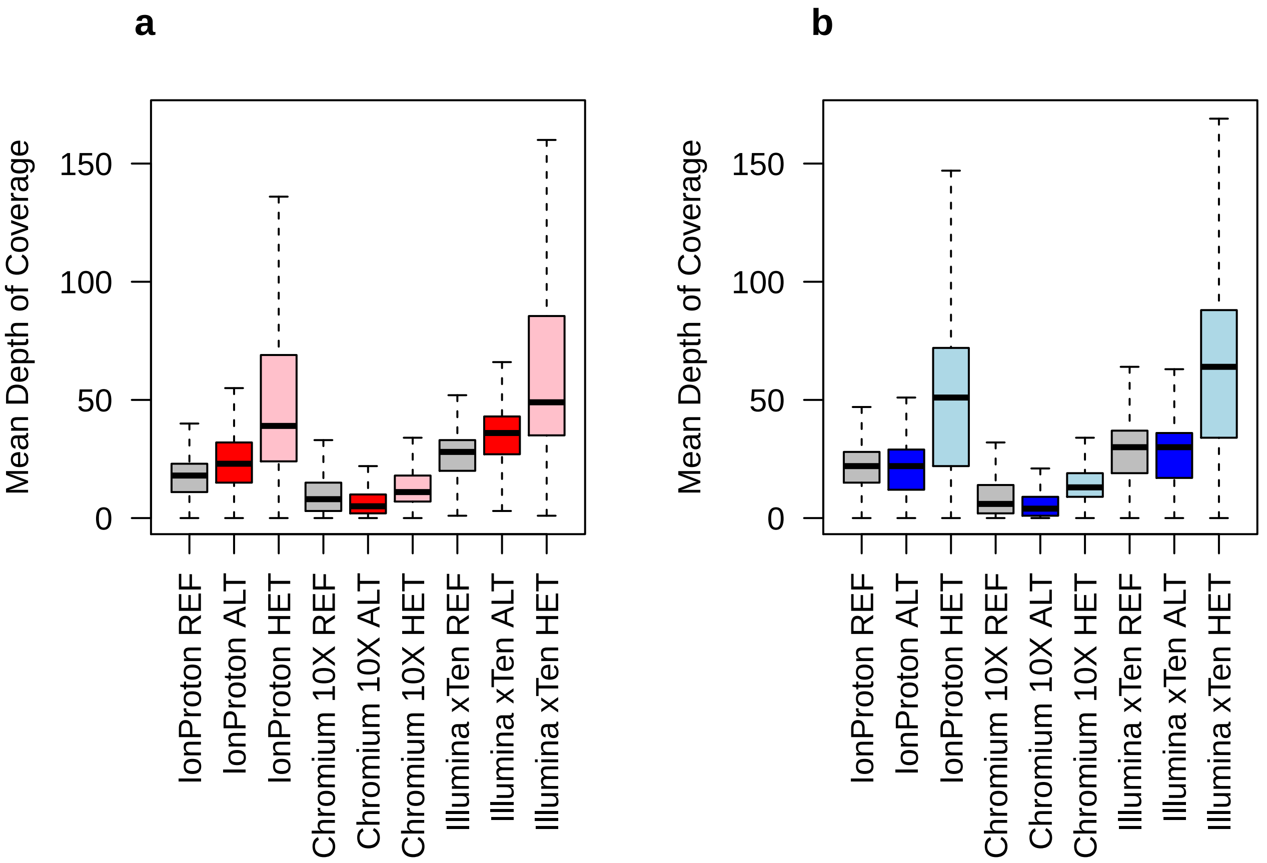
